# An NAD^+^–dependent sirtuin depropionylase and deacetylase (Sir2La) from the probiotic bacterium *Lactobacillus acidophilus* NCFM

**DOI:** 10.1101/252379

**Authors:** Sita V. Olesen, Nima Rajabi, Birte Svensson, Christian A. Olsen, Andreas S. Madsen

## Abstract

Sirtuins—a group of NAD^+^-dependent deacylases—have emerged as key in the connection between NAD^+^ metabolism and aging. This class of enzymes hydrolyze a range of *ε*-*N*-acyllysine PTMs and determining the repertoire of catalyzed deacylation reactions is of high importance to fully elucidate the roles of a given sirtuin. Here we have identified and produced two potential sirtuins from the probiotic bacterium *Lactobacillus acidophilus* NCFM and screening more than 80 different substrates, covering 26 acyl groups on five peptide scaffolds, showed that one of the investigated proteins—Sir2La—is a *bona fide* NAD^+^-dependent sirtuin, catalyzing hydrolysis of acetyl‐, propionyl‐, and butyryllysine. Further substantiating the identity as a sirtuin, known sirtuin inhibitors nicotinamide and suramin as well as a thioacetyllysine compound inhibit the deacylase activity in a concentration-dependent manner. Based on steady-state kinetics Sir2La showed a slight preference for propionyllysine over acetyllysine and butyryllysine, driven both by *K*_M_ (14 μM *vs* 21 μM and 15 μM) and *k*_cat_ (4.4·10^−3^ s^−1^ *vs* 2.5·10^−3^ s^−1^ and 1.21·10^−3^ s^−1^). Moreover, while NAD^+^ is a prerequisite for Sir2La-mediated deacylation, Sir2La has very high *K*_M_ for NAD^+^ compared to the expected levels of the dinucleotide in *L. acidophilus*. Sir2La is the first sirtuin from Lactobacillales and of the Gram-positive bacterial subclass of sirtuins to be functionally characterized. The ability to hydrolyze propionyl‐ and butyryllysine emphasizes the relevance of further exploring the role of other short-chain acyl moieties as PTMs.

---

Nicotinamide adenosine dinucleotide (NAD^+^) is well known for its role—together with the reduced form, NADH—as an essential redox couple in metabolism. While the dinucleotide is intact during oxidoreductase reactions, NAD^+^ also serves as substrate in reactions where the charged nicotinamide moiety acts as leaving group rather than hydride acceptor. NAD^+^-consuming enzymes include deacylases (sirtuins), cyclic adenosine diphosphate synthetases (cADPRS), and poly(adenosine diphosphate ribose) polymerases/adenosine diphosphate-ribosyl transferases (PARPs/ARTs). Dinucleotide turnover resulting from the activity of these enzymes leaves cells in constant need of NAD^+^-supply. In mammals, NAD^+^ is synthesized either via a *de novo* pathway from tryptophan, or via salvaging the three precursor vitamins nicotinic acid, nicotinamide, or nicotinamide riboside (NR). The recommended dietary allowance of vitamin B_3_ (niacin, a generic term for nicotinic acid and nicotinamide) is 15–20 mg/day, an amount sufficient to prevent clinical symptoms of niacin deficiency. However, in recent years significant interest has been raised around the use of NAD^+^-precursors as antiaging agents (1,2,3,4). Thus, NR is available as a commercial supplement and has been tested in clinical trials with intake up to 1 g/day (equivalent to 420 mg nicotinamide) (5,6). Oral administration of supplements makes these compounds available to the microbiota present in the gastrointestinal tract proximal to the area of absorption that occurs in both the stomach and small intestine (7). The endogenous gut microbiota is dominated by organisms from the Firmicutes, Bacteroidetes, Actinobacteria and Proteobacteria phyla (8), and is an increasingly recognized contributor to health and disease (9,10). A high intake of NAD^+^-precursors may impact both auto‐ and allochthonous commensal bacteria and emphasizes the relevance of characterization of microbial enzymes involved in NAD^+^-metabolism (11). *L. acidophilus* is a Gram-positive, lactic acid-producing firmicute used extensively as a probiotic in foods and dietary supplements (12), notably motivating further investigation.

In particular the sirtuin class of enzymes have emerged as key in the connection between NAD^+^ metabolism and aging (13,14). Named after the yeast enzyme *silent information regulator 2* (Sir2), originally shown to regulate mating type genes (15,16,17), sirtuins and their role in aging have been extensively investigated. Initially, *Saccharomyces cerevisiae sir2*-mutants were found to exhibit decreased lifespan, whereas an additional copy of the *sir2* gene conferred extended lifespan (18). Similarly sirtuin-mediated longevity has been shown in fruit flies (19), nematodes (20), and mice (21). The canonical function of sirtuins utilizes NAD^+^ as co-substrate to catalyze hydrolysis of *ε*-*N*-acyllysine, releasing nicotinamide and transferring the acyl moiety to form 2′-*O*-acyl adenosine diphosphate ribose (e.g., 2′-*O*-acetyl adenosine diphosphate ribose (OAcADPR)). Lysine acetylation is a prevalent, reversible PTM associated with a wide range of biological processes within all domains of life (22). Sirtuins are involved in regulation of gene expression due to their role in controlling the acetylation pattern on histones. Lysine acetylation can be reversed by two classes of enzymes, the zinc-dependent histone deacetylases (HDACs) and the sirtuins. Even though all sirtuins harbor a conserved catalytic core domain (23,24) phylogenetic analysis reveals distinct classes (24) and it has become apparent that the substrate specificity of sirtuins varies with these classes. Eukaryotic sirtuins are primarily found in classes I-IV, archaeal sirtuins belong to classes III and U, whereas bacterial sirtuins are found in classes II, III, and U (25), as well as in the class of Sir2-like enzymes (Fig. 1a). Additionally, a novel class M comprises bacterial and eukaryotic enzymes (26). While sirtuins in class U (abbreviation for “undifferentiated”) were originally grouped as a single class, it is now realized to comprise at least four sub-classes (25). Thus, in the NCBI Conserved Domain Database (CDD) (27), two families that fall into the class U are defined; one comprising both archaeal and bacterial enzymes based on homology with *Archaeoglobus fulgidus* Sir2Af2 [NCBI CDD cd01413] and one comprising sirtuins from Gram-positive bacterial species and fusobacteria [NCBI CDD cd01411].

**Figure 1.**
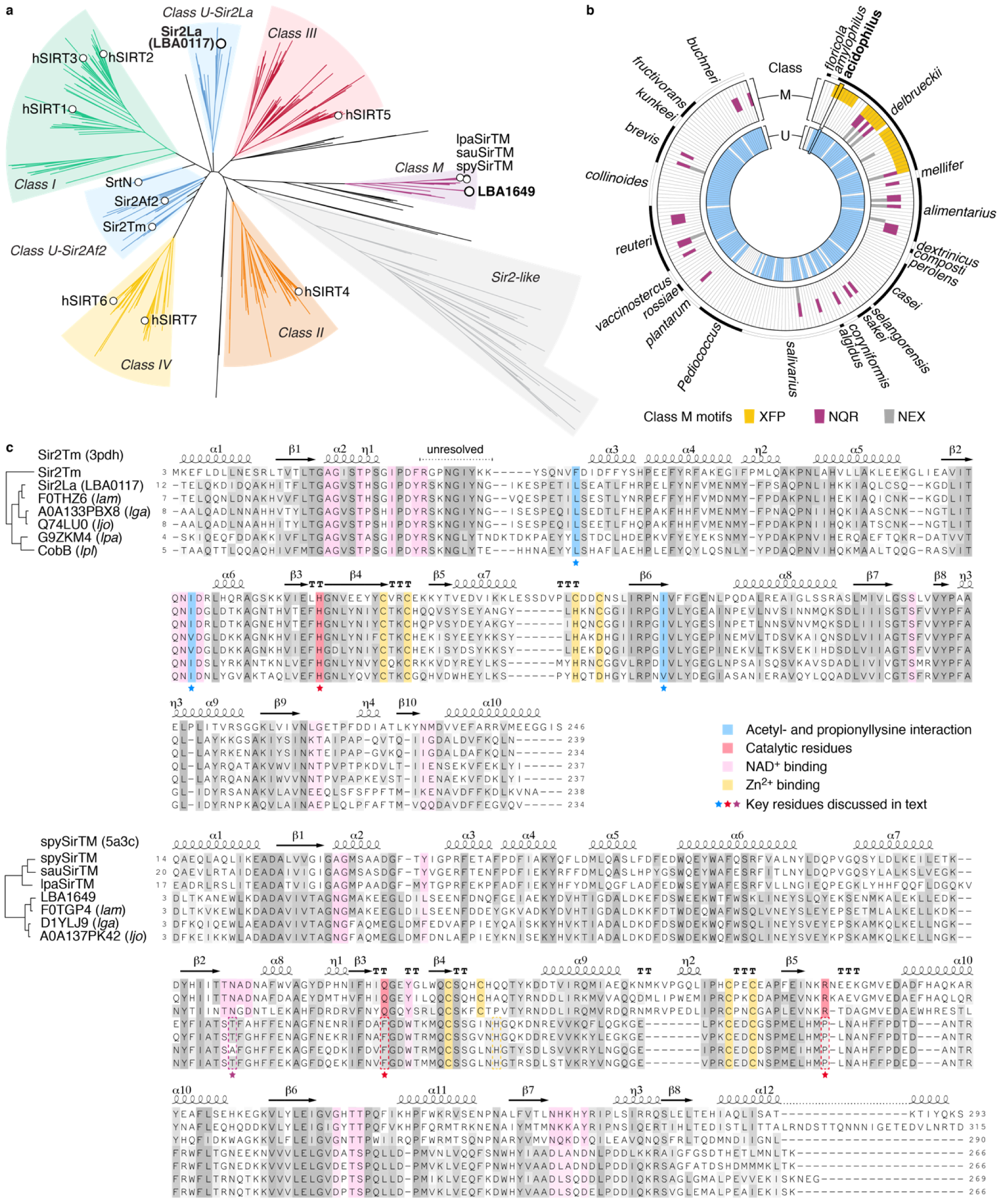
Bioinformatic analysis. a) Phylogenetic tree of 535 sirtuins with indication of sirtuin classes. The positions of the two putative sirtuins from *L. acidophilus* NCFM (in bold), the human sirtuins 1–7, as well as selected sirtuins from class U and M are indicated. b) Sirtuins in *lactobacilli*. BLAST searches of classes U and M sirtuins in lactobacilli. Three different motifs of three functionally critical residues are identified in the class M hits. c) Alignment of selected sirtuins. Secondary structure assignment of known crystal structures and key residues discussed in the text are shown.

Although less prevalent than lysine acetylation (28), a range of other acyl groups have been identified as PTMs on histones and proteins in general through proteomics studies. These modifications include the short-chain acyl groups formyl (29,30), propionyl (31,32), butyryl (31) and crotonyl (33), the carboxyacyl groups malonyl (34), succinyl (35) and glutaryl (36), as well as the long-chain aliphatic group myristoyl (37,38). Of the seven identified human sirtuins, only hSIRT1–3 are efficient deacetylases in *in vitro* studies, whereas hSIRT6-catalyzed deacetylation can be achieved by addition of fatty acids to the reaction mixture (39) and hSIRT6 is also a functional deacetylase activity *in vivo* (40). Hydrolysis of long-chain acyl PTMs is catalyzed by hSIRT1–3 (39) and 6 (41), hSIRT5 has been shown to possess lysine demalonylase, desuccinylase, and deglutarylase activity (36,42), and hSIRT4 was recently demonstrated to catalyze cleavage of lipoyl‐ and biotinyllysine (43) as well as glutaryl-, 3-methylglutaryl‐, 3-hydroxy-3-methylglutaryl‐ and 3-methylglutaconyllysine (44,45). Some sirtuins act on several types of acyllysines as substrates. Most prominent is the ability of class I sirtuins to hydrolyze both acetyl-and long-chain acyllysines, but deacylation of propionylated and butyrylated peptides and proteins have also been described by hSIRT1–3, the yeast sirtuin HST2, CobB from *Salmonella enterica*, and Sir2 from *Thermotoga maritima* (Sir2Tm) (46,47,48). The sirtuin from *E. coli*, CobB, has also been demonstrated to harbor lysine deacetylase, desuccinylase, and delipoamidase activity (49,50,51). Although sirtuins are primarily recognized as deacylases, hSIRT4 (52), hSIRT6 (53,54,55), and other sirtuins (56,57,58) were reported to be ADP-ribosyl transferases. Similarly, a new recently described distinct class of sirtuins (class M or SirTMs) was identified primarily in a range of pathogenic microorganisms including both bacteria and fungi (26). Notably, class M sirtuins are devoid of deacylase activity but rather catalyze a specific ADP-ribosylation. The modified protein GcvH-L (glycine cleavage system H-like) belongs to the same extended operon as SirTM, and GcvH-L is only ADP-ribosylated when lipoylated—a PTM installed by LplA2 (lipoate protein ligase A), a third protein in the same extended operon (26). Thorough characterization of substrate specificity of any sirtuin is therefore of high importance to fully elucidate the various roles in the cell.

While many sirtuins across a range of species have been investigated on both a molecular, cellular and organismal level, characterization of enzymatic activity of any sirtuin from the Lactobacillales order is still missing. Motivated by the potential impact of NAD^+^-precursor supplements on the gut microbiome, probiotic use of *L. acidophilus*, and relevance of sirtuins in gene regulation and general cell maintenance, we analyzed the *L. acidophilus* NCFM genome and cloned and produced the two identified putative sirtuins. This enabled characterization of substrate scope, enzymatic efficiency and inhibition by known and novel sirtuin inhibitors of what is to our knowledge the first NAD^+^-dependent deacylase from a Lactobacillales species.

## Results

### Identifying sirtuins in *L. acidophilus* NCFM

Bioinformatics analysis revealed two putative sirtuins (LBA0117 [UniProt Q5FMQ6] (which we will refer to as Sir2La) and LBA1649 [UniProt Q5FIL3]) in the genome sequence of *L. acidophilus* NCFM (59). Multiple sequence alignment and phylogenetic analysis placed these two proteins in the U and M classes of sirtuins, respectively (Fig. 1a and 1c, Tables S1–S3). BLAST searches identified similar proteins in many lactobacilli across the genus (Fig. 1b and Table S4). To verify their function as NAD^+^-dependent deacylases and characterize the potential substrate scope the two proteins were produced recombinantly in *E. coli* and profiled using a fluorogenic assay including 26 acyl modifications on five peptide scaffolds with NAD^+^ as co-substrate. The peptide sequences were based on known lysine acylation sites in histone 3 (H3, K9), histone 4 (H4, K12), tumor suppressor p53 (p53, K320), and dihydrolipoyllysine acetyltransferase (DLAT, K259). Linear acyl groups ranging from one to fourteen carbon atoms in length, branched chain, unsaturated, hydroxylated, and carboxyacyl groups were included (Fig. 2 and S1). With NAD^+^ as co-substrate Sir2La was found to efficiently deacetylate, depropionylate, and debutyrylate modified lysine residues, thereby unequivocally establishing the enzyme as a *bona fide* sirtuin. In line with the reported function of class M sirtuins as ADP-ribosyl transferases, LBA1649 had no measurable deacylase activity under the given conditions of the 15 tested acyl groups.

**Figure 2.**
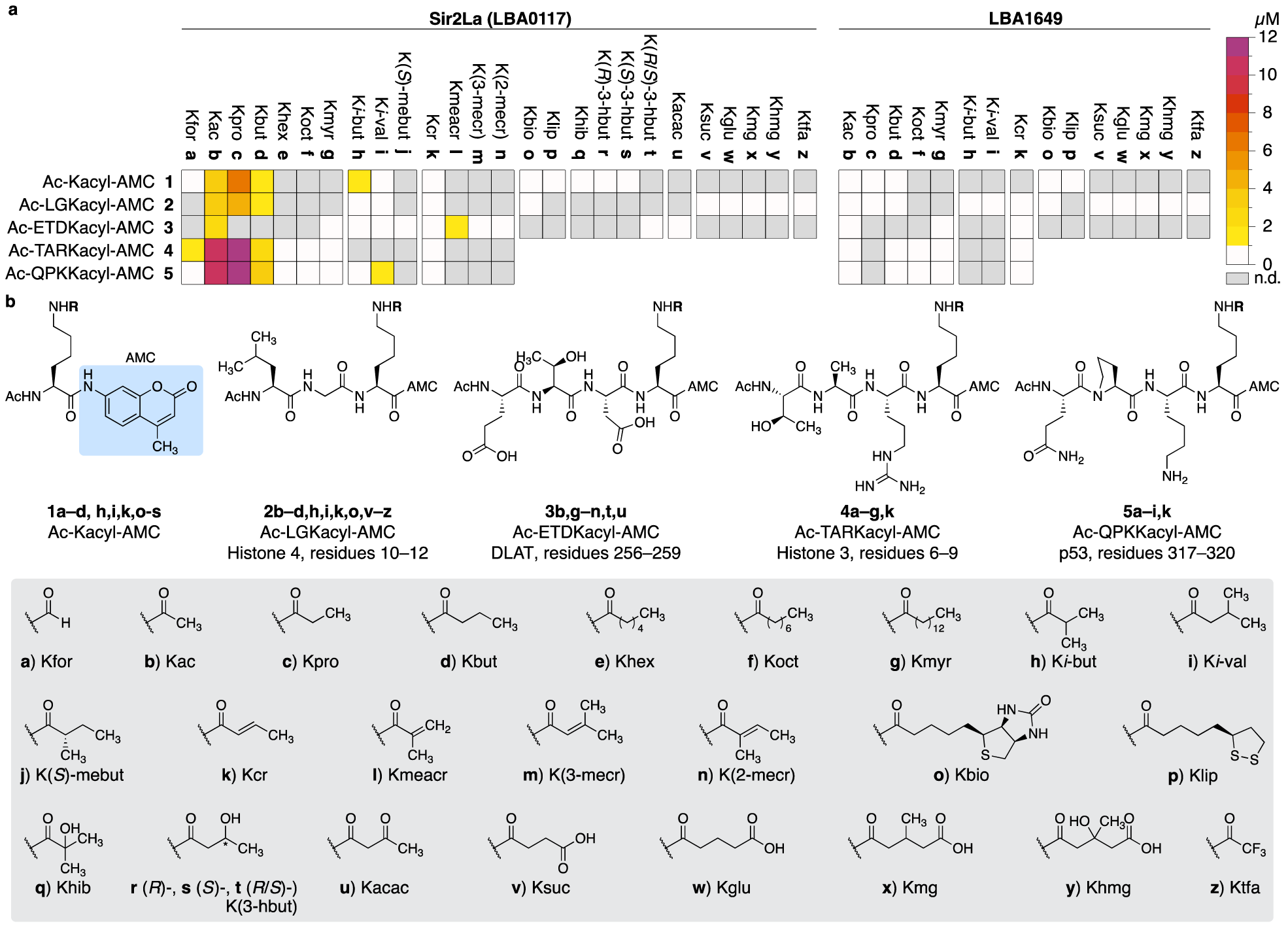
Substrate screening of Sir2La and LBA1649. a) Heatmap of Sir2La and LBA1649-dependent deacylase activity based on endpoint assays. All reactions were incubated for 1 h at 37 °C before development and fluorescence analysis. b) Substrate structures; amino acid sequences and acyl groups. n.d., not determined.

### Determining Michaelis-Menten parameters

Having established the overall substrate scope of Sir2La, the hydrolase activity was further characterized against acetyl, propionyl, and butyryl substrates and investigated with a range of sirtuin inhibitors, in assays that allow continuous monitoring of product formation. The initial screen indicated that neighboring residues carrying a positively charged side chain promoted enzyme activity (series **4** and **5**, Fig. 2). However, the use of trypsin as developer in the continuous assay precludes unmodified lysine and arginine residues, since both the substrate and product can be hydrolyzed C-terminal to these residues by trypsin leading to altered affinity and kinetics (60). The use of histone 4 lysine 12 derived substrates (series **2**, Fig. 2) is well-established and substrates containing Kac (*ε*-*N*-acetyllysine) (**2b**), Kpro (*ε*-*N*-propionyllysine) (**2c**), and Kbut (*ε*-*N*-butyryllysine) (**2d**) were chosen for kinetic and inhibition studies. Under steady-state conditions, the initial rates of hydrolase activity at varying peptide substrate and constant [NAD^+^] (Fig. 3a) were used to obtain the Michaelis-Menten constant (*K*_M_) for Sir2La-dependent deacylation (Table 1). Lower affinity was found for the Kac than the Kpro and Kbut substrates and *k*_cat_ against the Kpro substrate was twice that of the Kac substrate, which in turn yielded higher *k*_cat_ than the Kbut substrate. This substrate preference is reflected in *k*_cat_/*K*_M_ indicating that Sir2La-dependent deacylation is most efficient for depropionylation (320 ± 90 M^−1^s^−1^), compared to deacetylation (120 ± 40 M^−1^s^−1^), and debutyrylation (82 ± 17 M^−1^s^−1^) (Table 1). The values are comparable or slightly lower than previously reported for deacylation reactions by sirtuins (41,42,45,46,60,61).

**Figure 3.**
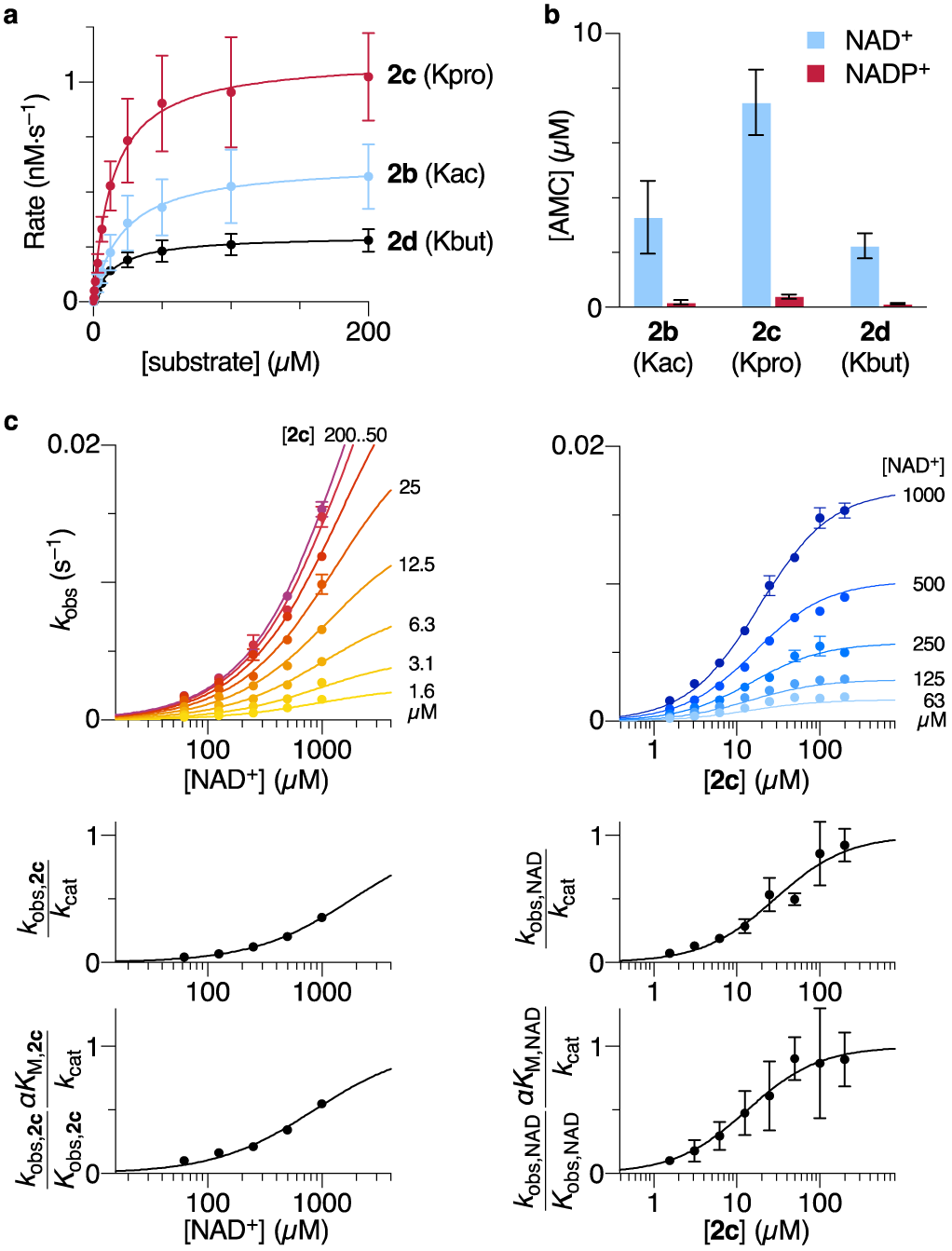
Characterization of deacylation reactions by Sir2La. a) Michaelis-Menten plots for Sir2La-dependent deacylation of three short-chain substrates measured at 500 μM NAD^+^. b) Co-substrate selectivity for Sir2La. Bar-graphs of substrate deacylation using either NAD^+^ (blue) or NADP^+^ (red). c) Bisubstrate kinetic measurements for Sir2La-dependent deacylation of *ε*-*N*-propionyllysine substrate **2c**, showing the global fit of steady-state rates to the sequential random order equation and the corresponding secondary plots of (*k*_obs_/*k*_cat_) and (*k*_obs_/*K*_obs_·*αK*_M_/*k*_cat_) as functions of substrate concentrations. Best fit is shown as lines. All reactions were monitored for at least 1 h at 25 °C.

**Table 1.** Kinetic values for Sir2La-catalyzed deacylation of *ε*-*N*-acyllysine substrates **2b** (Kac substrate), **2c** (Kpro substrate), and **2d** (Kbut substrate) measured at 500 μM NAD^+^.

| | $K_M$<br>($\mu\text{M}$) | $k_{\text{cat}}$<br>($\cdot 10^{-3} \text{ s}^{-1}$) | $k_{\text{cat}} \cdot K_M^{-1}$<br>($\text{M}^{-1} \text{ s}^{-1}$) |
| --- | --- | --- | --- |
| Ac-LGKac-AMC (**2b**) | $21 \pm 6$ | $2.5 \pm 0.2$ | $120 \pm 40$ |
| Ac-LGKpro-AMC (**2c**) | $14 \pm 3$ | $4.4 \pm 0.3$ | $320 \pm 90$ |
| Ac-LGKbut-AMC (**2d**) | $15 \pm 2$ | $1.21 \pm 0.06$ | $82 \pm 17$ |

The NAD^+^-dependency on deacylation rate also prompted testing of NADP^+^ as a possible co-substrate for Sir2La-dependent deacylation, but incubation with NADP^+^ had only minor effect on deacylation activity (Fig. 3b).

Since the propionyllysine substrate was the most efficiently converted, bisubstrate kinetics of Sir2La-catalyzed depropionylation was determined by varying both peptide substrate and co-substrate concentrations (Fig. 3c). In agreement with the general mechanism of sirtuins, the data supports a sequential mechanism and not a ping-pong mechanism, i.e., both peptide‐ and co-substrate are required to bind for catalysis to occur. The data however, do not support an ordered, rapid-equilibrium mechanism (Fig. S2), but rather either an ordered steady-state mechanism (i.e., turnover of the ternary complex is not the rate-limiting step) or a mechanism with random binding of peptide‐ and co-substrate (kinetic values are shown in Table 2). These two mechanisms cannot be distinguished based on the present experiments.

**Table 2.** Kinetic values derived from bisubstrate kinetic measurement of Sir2La-catalyzed deacylation of Kpro substrate **2c**.

| | $k_1$<br>($\mu\text{M}^{-1} \text{ s}^{-1}$) | $k_{-1}$<br>($\text{s}^{-1}$) | $K_{M,2c}$<br>($\mu\text{M}$) | $K_{M,\text{NAD}} / K_{M,\text{NAD,app}}$<br>($\mu\text{M}$) | $\alpha$ | $k_{\text{cat}} / k_3$<br>($\cdot 10^{-3} \text{ s}^{-1}$) |
| --- | --- | --- | --- | --- | --- | --- |
| Random binding, rapid equilibrium | — | — | $13 \pm 2$ | $860 \pm 250$ | $2.1 \pm 0.8$ | $48 \pm 4$ |
| Ordered binding, steady state | $1.7 \pm 0.7$ | $22 \pm 9$ | $13 \pm 2$ | $1850 \pm 880$ | — | $48 \pm 4$ |

### Inhibition of Sir2La hydrolase activity

The efficiency of a range of inhibitors originally developed for human sirtuins was tested on the Sir2La-dependent deacylation activity (Fig. 4). Notably, only a few of these compounds exhibited significant inhibitory activity (Fig. 4b, Table 3 and Fig. S3). The pan-sirtuin inhibitors suramin and nicotinamide reduced the deacylation reaction rates at high micromolar concentrations (Table 3), whereas sirtinol (hSIRT2, IC_50_ = 38 μM) (62), AGK2 (hSIRT2, IC_50_ = 3.5 μM) (63), and SirReal2 (hSIRT2, IC_50_ = 0.4 μM) (64) at concentrations up to 100 μM failed to inhibit the activity (Fig. S3). Thioacyllysine-based inhibitors are validated sirtuin inhibitors and gratifyingly, the recently reported thioacetamide **6** (65) is here found to inhibit Sir2La, whereas a propionyllysine-inspired thiourea (**7**) was a poor inhibitor (77% residual Sir2La-activity at 50 μM using **2c** as substrate). Adenosine diphosphate ribose (ADPR), NADH, and NADPH were tested as potential inhibitors but in agreement with previous results for human and yeast sirtuins (60,66) no significant inhibition was found below 10 mM (Fig. 4b and Fig. S3).

**Figure 4.**
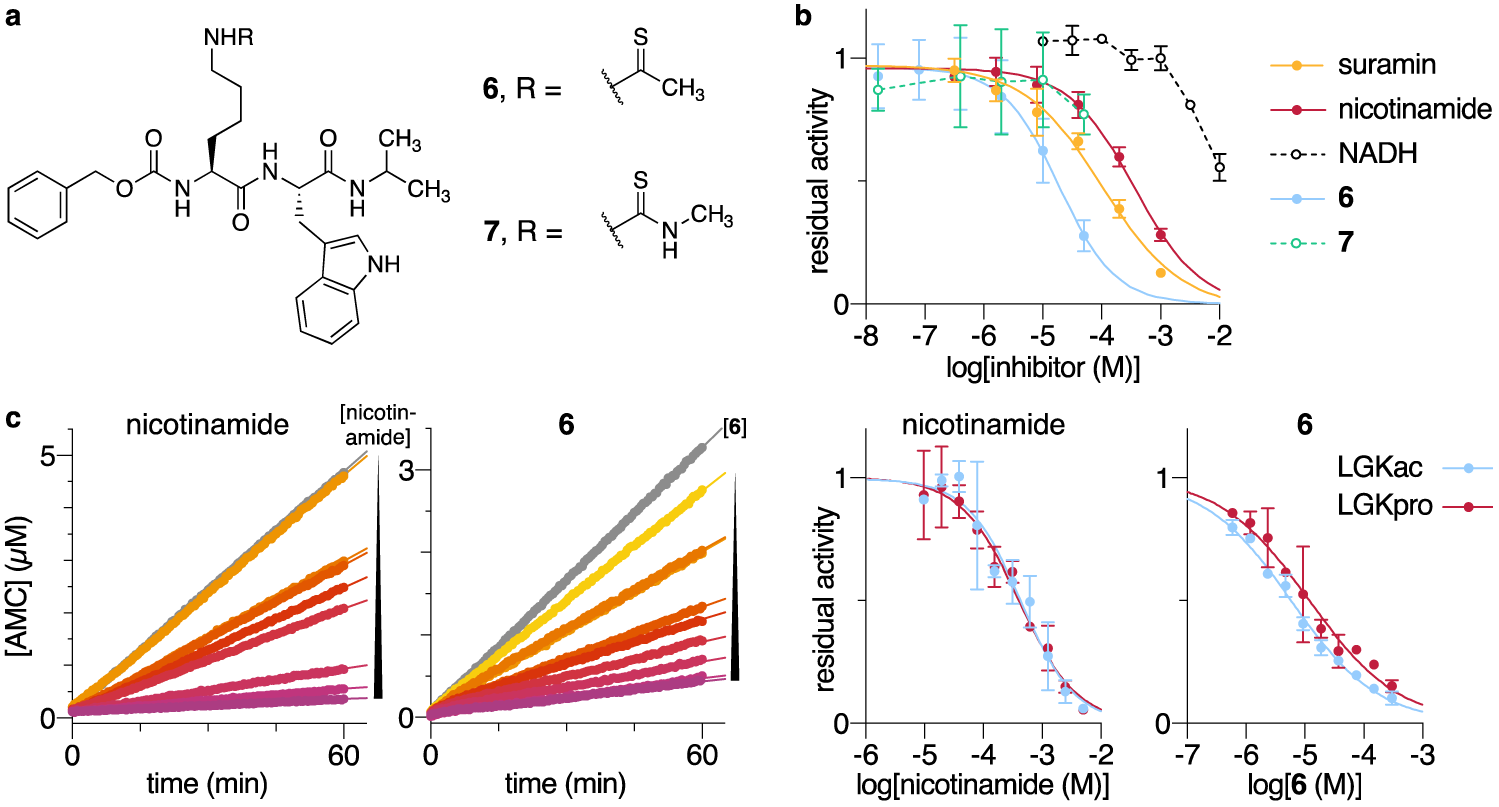
Inhibition of Sir2La-catalyzed deacylation of Kpro substrate **2c**. a) Structures of inhibitors **6** and **7**. b) Residual activities for depropionylation, dose-response curves from end-point assays. c) Progression curves and data fitting for inhibition of deacylation by nicotinamide and compound **6**.

**Table 3.** IC50-values for inhibition of Sir2La-catalyzed deacylation, measured in end-point assays.

| Substrates | Inhibitors ($\text{IC}_{50}$ ($\mu\text{M}$)) | | |
| --- | --- | --- | --- |
|  | nicotinamide | suramin | **6** |
| Ac-LGKac-AMC (**2b**) | $460 \pm 20$ | $110 \pm 20$ | $16.1 \pm 0.2$ |
| Ac-LGKpro-AMC (**2c**) | $350 \pm 70$ | $100 \pm 30$ | $15 \pm 2$ |
| Ac-LGKbut-AMC (**2d**) | $51 \pm 31$ | $640 \pm 370$ | $20 \pm 5$ |

To determine the kinetics of inhibition by nicotinamide and thioacetamide **6**, initial velocities of Sir2La-mediated deacylation of substrates **2b** and **2c** were measured at varying inhibitor concentrations. Maintenance of a steady-state rate over 60 min demonstrated that both compounds inhibit deacylation via a fast-on–fast-off mechanism. (Fig. 4c and Table 4).

**Table 4.** IC_50_-values for inhibition of Sir2La-catalyzed deacylation, measured in continuous assays.

| Substrates | Inhibitors ($\text{IC}_{50}$ ($\mu\text{M}$)) | |
| --- | --- | --- |
|  | nicotinamide | **6** |
| Ac-LGKac-AMC (**2b**) | $460 \pm 240$ | $5.8 \pm 0.1$ |
| Ac-LGKpro-AMC (**2c**) | $400 \pm 20$ | $12 \pm 4$ |

## Discussion

The first sirtuin (Sir2La) from a species belonging to the Lactobacillales order has been identified, produced recombinantly and functionally characterized. This and a second candidate sirtuin (LBA1649) were tracked by genome mining of *L. acidophilus* NCFM. Sir2La efficiently catalyzes deacylation of propionyl-, acetyl-, and butyryllysine substrates in an NAD^+^-dependent manner. Additionally, the enzyme activity was inhibited by known sirtuin inhibitors and enzyme kinetics revealed a sequential binding of the two substrates following either a random or steady-state ordered mechanism.

Phylogenetic analysis categorizes Sir2La and LBA1649 as class U and M sirtuins, respectively (Fig. 1a), and similar proteins are encoded by other lactobacilli (Fig. 1b), including closely related *L. amylovorus*, *L. gasseri*, and *L. johnsonii*—all four species belonging to the socalled *L. delbrueckii*-group—and more distantly related *L. plantarum* and *L. parafarraginis* (67,68) (Fig. 1c). Based on phylogeny, LBA1649 belongs to class M sirtuins and as such failed to deacylate any of the investigated substrates. Notably, while Ahel and coworkers found that *L. parafarraginis* express a class M sirtuin in the operon also encoding GcvH-L and LplA2, they did not expand the investigation of the genus (26). Sequence alignment of LBA1649 with SirTM from *L. parafarragins* and prototype class M sirtuins spySirTM [UniProt P0DN71] and sauSirTM [UniProt A0A0H3JQ59] highlighted mutations of three functionally critical residues. A glutamine resulting from the mutation of a histidine normally involved in deacylation (e.g., hSirt2-H187 or Sir2La-H124), an arginine involved in correct positioning of NAD^+^ in the active site, and a nearby asparagine (Q137, R192, and N118 of spySirTM, respectively), were identified as critical for the catalytic ADP-ribosyltransferase activity of SirTM enzymes. While these residues are indeed conserved in *L. parafarragins*, corresponding residues for LBA1649 and closely related homologs from *L. johnsonii*, *L. amylovorus* and *L. gasseri* are phenylalanine (F127), proline (P175), and threonine (T108) or alanine, respectively (Fig. 1c). Based on the proposed model of SirTM-mediated ADP-ribosylation these residues are not expected to partake in catalysis of neither deacylation nor ADP-ribosylation. Interestingly, a BLAST search of lactobacilli proteins in the NCBI database, revealed that LBA1649-like proteins (containing the XFP motif, X = A/S/T/Y) were found exclusively in the *L. delbrueckii* group, whereas proteins containing the catalytically competent NQR motif were represented in the full lactobacilli phylogenetic tree including the *L. delbrueckii* group (Fig. 1b and Table S4). We do not at present have a hypothesis of the function of LBA1649 (and homologous proteins) and while resolving the function of LBA1649 would be relevant, we decided not to pursue this further, hence proceeding only with Sir2La.

Of the subclass of sirtuins from Gram-positive bacterial species and fusobacteria (Class U-Sir2La, Fig. 1a), Sir2La appears to be the first functionally characterized enzyme. Previously characterized class U sirtuins include Sir2Af2 from *A. fulgidus*, Sir2Tm from *T. maritima*, and SrtN from *B. subtilis* (51,69), all belonging to the class U subclass with Sir2Af2-homology (Class U-Sir2Af2, Fig. 1a). In the present study, only deacetylation, depropionylation and debutyrylation were efficiently catalyzed by Sir2La (Fig. 2). Similar to previous findings for human sirtuins (60,70,71), deacylation efficiency depended to some extent on the peptide scaffold. However, Sir2La failed to hydrolyze substrates containing acyl groups with chains longer than four carbon atoms including the dithiolane-containing lipoyl group. Such strict short-chain acyl substrate requirement for Sir2La has also been reported for Sir2Tm (50), but contrasts the substrate specificity for other investigated sirtuins, including cleavage of short- and long-chain acyl groups by human sirtuins 1–3 (39,60), cleavage of acetyl and lipoyl by SrtN from *B. subtilis* (51,69), as well as the very broad substrate scope of CobB from *E. coli* (49,50,51). Archaeal Sir2Af2 can bind and hydrolyze acetylated, propionylated, and butyrylated as well as myristoylated peptides with increasing efficiency (50). This highlights a differential substrate preference for the class U subclasses. Sir2Tm has been crystallized with acetylated [PDB ID 2h2d] and propionylated [PDB ID 3pdh] peptides but not with long-chain acylated (e.g., myristoylated) peptides. On the other hand, reflecting substrate preference Sir2Af2 has been crystallized with a myristoylated peptide [PDB ID 4ywj] bound in the active site, which revealed a hydrophobic pocket similar to the one found in class I sirtuins. The experimental evidence suggests that the structure of Sir2La would resemble that of Sir2Tm, and sequence alignment indicated an important overlap between the two proteins (32% identity and 49% similarity, Table S2), including similar residues flanking the binding pocket of the acyl groups in Sir2Tm (Fig. 1c).

Sir2Tm deacetylates and depropionylates with similar efficiency whereas debutyrylation is less efficient (46,50). Similar to Sir2La, a slightly lower *K*_M_ for propionylated vs acetylated substrates (14 mM vs 21 mM) was reported for both Sir2Tm (87 mM vs 104 mM) (46) and yHST2 (8.6 mM vs 21 mM) (47,72), however, *k*_cat_/*K*_M_ for Sir2Tm-mediated deacylation of the two substrates was 1.2·10^3^ M^−1^s^−1^ and 1.4·10^3^ M^−1^s^−1^, respectively. Similar efficiency of deacylation has been proposed to stem from the slower release of 2′-*O*-propionyl adenosine diphosphate ribose (OPrADPR) than of OAcADPR, likely as a result of the greater hydrophobicity of the former (46). For Sir2La, the same reasoning would favor deacetylation and the origin of this reversed substrate preference has yet to be established.

The present study also shows that NAD^+^ is necessary for Sir2La-dependent deacylation reactions to take place, which further strengthens the conclusion that Sir2La is a deacylase of the sirtuin-type. Previous investigations found *K*_M_ values for NAD^+^ to vary significantly among class I sirtuins. yHST2 appears to have remarkably high affinity for NAD^+^ (*K*_M_ of 2 μM was measured applying an acetylated peptide (73)) whereas *K*_M_ ranging from 18 to 360 μM was reported for other enzymes (47,57,66,73,74). An even higher *K*_M_ of >800 μM was found for Sir2La, highlighting a potential difference between sirtuin classes. In eukaryotes, the total intracellular dinucleotide concentrations are reported to 1–3 mM (75), free NAD^+^ concentrations being estimated to 100 μM in the cytosol and nucleus and 230 μM in mitochondria (76), placing the reported *K*_M_ values in a physiologically relevant range (77). Investigation of total NAD^+^ concentrations in prokaryotes dates back to the 1960’s and 1970’s. *L. acidophilus* NCFM is a microaerophile—exposed to low levels of oxygen, and a correlation between aerobic requirement and NAD^+^ content has been suggested (78) with obligate anaerobes having a concentration similar to yeast (>1 mM) and lower concentrations for facultative anaerobes (1–0.2 mM) and obligate aerobes (<0.2 mM) (79). However, in a more recent study using ^13^C and ^31^P NMR, total concentrations in the facultative anaerobic bacterium *Lactococcus lactis* were investigated (80) and under both anaerobic and aerobic conditions, [NAD^+^] was ∼5 mM, while NADH was undetectable (<0.5 mM). Thus, the total [NAD^+^] ranges from 0.2–5 mM, however, we have not been able to find estimates of free concentrations of the relevant dinucleotides in bacteria. The relation between NAD^+^ concentration and *K*_M_ value will impact sirtuin activity, but the potential relevance of the observed higher *K*_M_ value for Sir2La is uncertain.

In agreement with the established mechanism for sirtuin deacylation our data support a sequential mechanism – i.e., both peptide substrate and NAD^+^ bind before the reaction proceeds. Additionally, the hyperbolic secondary plots (Fig. 3b) indicate a steady-state ordered mechanism or a random binding order for the two substrates. Sirtuins from classes I, III, and U (Sir2Af2) have been shown to follow an ordered mechanism, where the proper NAD^+^-binding pocket is formed only after binding of the peptide substrate (73,74,81,82). Sirtuin 6 (class IV) is the only investigated enzyme that appears to follow a random binding order (74,82). Further experiments are needed to determine whether the reaction of Sir2La follows a random substrate binding or steady-state ordered mechanism and further compare to previously investigated sirtuins.

To further substantiate that the Sir2La-deacylation mechanism is analogous to that of other sirtuins, the effect of known sirtuin inhibitors was analysed. This included suramin, sirtinol, AGK2, and SirReal2, the NAD^+^-derivatives nicotinamide, ADPR, NADH, and NADPH as well as substrate-derived compounds **6** and **7**. In agreement with literature (60,66), where NADH has been reported to inhibit some human sirtuins at high concentrations, millimolar concentrations of NADH and ADPR were needed to inhibit Sir2La-catalyzed depropionylation and deacetylation. On the other hand, nicotinamide, suramin, and thioacetamide **6** inhibited deacylation at micromolar concentrations (Fig. 4 and Tables 3 and 4). Noticeably, the potency of nicotinamide was higher for Sir2La-dependent debutyrylation than for deacetylation and depropionylation (IC_50_ of 50 μM, 460 μM and 350 μM, respectively). The two substrate-derived compounds, thioacetamide **6** (65) and thiourea **7** (Fig. 4a), contain chemotypes that form stalled intermediates in the active site of sirtuins (65,83,84,85,86). In accordance with previous strong inhibition of sirtuins by thioamide-based inhibitors, thioacetamide **6** was the most potent of the present investigated inhibitors. The peptide scaffold of compounds **6** and **7** was developed for efficient inhibition of human sirtuin 5 through an iterative process including optimization of residues adjacent to the modified lysine (65) and improved inhibition of Sir2La-activity may be expected through a similar Sir2La-targeted process. Surprisingly, even though Sir2La shows a slight preference for propionylated substrates, thiourea **7** did not significantly decrease deacylase activity on any of the three substrates at the tested concentrations. We hypothesize that the planar geometry imparted by the thiourea moiety led to a relatively rigid acyl-surrogate that may prevent the inhibitor to adopt a suitable orientation within the active site and to react with NAD^+^.

The cellular relevance of lysine propionylation and also butyrylation is increasingly being investigated. Propionyl‐ and butyryllysine residues were initially identified on histones in human (31,87) followed by yeast cell lines (88). Subsequently, these PTMs have been described either as competing with other PTMs (e.g., Kbut vs Kac) or combined with other modifications (89,90). Similar to other acyl PTMs, identification of propionylation and butyrylation sites has been expanded to include proteins other than histones. Several acylation sites were identified on tumor suppressor p53 and histone acetyl transferases p300 and CBP in human non-small cell lung carcinoma cells (32), and elevated levels of the corresponding CoAs either as a result of impaired propionyl-CoA carboxylase or short-chain acyl-CoA dehydrogenase activity (91) or by chronic ethanol ingestion (92) causing increased acylation levels of both nuclear and mitochondrial proteins. Demonstrating a regulatory function of a site-specific modification, propionylation of propionyl-CoA synthetase at lysine-592 in *S. enterica* has been shown to inhibit its enzymatic activity, and interestingly the activity can be rescued by sirtuin-mediated depropionylation (48). Lysine acylation of other proteins that are either directly involved in or metabolically close up‐ or downstream to the generation of a reactive compound, e.g., an acyl-CoA or acyl phosphate, has also been reported (44,93). Furthermore, low-level acylation on thousands of sites has been reported throughout the proteome in *E. coli*, possibly formed through non-enzymatic acylation reactions (94). A critical component was shown to be acetyl phosphate, formed as an intermediate between acetyl-CoA and acetate. Deletion of CobB—the only *E. coli* sirtuin—lead to increased acetylation of a subset of these sites, primarily in unstructured regions and protein termini. Similarly, bacteria can convert between propionyl-CoA and propionyl phosphate and either may participate in the formation of propionyllysine throughout the proteome. Efforts to map bacterial lysine propionylomes have been reported (95,96,97) and suggest, as for lysine deacetylation, a link between lysine propionylation and metabolism. While the observed difference of *k*_cat_/*K*_M_ is only ∼2.7-fold between depropionylation and deacetylation, the favored depropionylation is intriguing, and a similar propionylome of *L. acidophilus* would be of high interest to further pursue the biological relevance of this observed preference.

With the importance of commensal bacteria in health and disease, a thorough characterization of microbial enzymes involved in NAD^+^-metabolism is important to understand both their primary role in the metabolism of the microorganism as well as predicting the potential impact of NAD^+^-precursor supplements on host-microbe interactions. In the present study we have identified, produced and characterized the first sirtuin from a species of the Lactobacillales order. The observed NAD^+^-dependent deacylation of acyllysine substrates unequivocally identifies Sir2La as a functional sirtuin enzyme, which is corroborated by its inhibition by nicotinamide, suramin and thioacetamide **6**. The observed ability to deacylate acetyl-, propionyl-, and butyryllysine residues emphasizes the relevance of further exploring the role of other short-chain acyl moieties than acetyl as PTMs, to further investigate the propionylome of microorganisms, and highlights a potential difference between the different subsclasses for class U sirtuins.

## Experimental procedures

### Identification of Sir2La

Using the Microbesonline (microbesonline.org) and UniProt (uniprot.org) databases and the COG0846 motif (98) as probe, two proteins (Sir2La/LBA0117 [UniProt Q5FMQ6] and LBA1649 [UniProt Q5FIL3]) were identified as putative lysine deacylases in the genome of *Lactobacillus acidophilus* NCFM (59).

### Multiple sequence alignment, phylogenetic tree

Based on Frye (24), Greiss and Gartner (25), Rack et al. (26), Okanishi et al. (96), and the SIR2 superfamily from the NCBI conserved domain database [NCBI CDD # cd00296] a list of 159 species was generated (Table S3). Subsequently, proteins assigned with sirtuin sequence motifs [eggNOG: COG0846, Pfam: pf02146, or PROSITE: ps50305] within these species were identified in the UniProt database, and the sequences truncated to comprise sirtuin domains identified in either UniProt or NCBI CDD. The resulting list of 1587 non-redundant domain sequences was curated by removal of closely related intraspecies sequences resulting in a list of 535 representative proteins (Table S3). One hundred alignments of these proteins were calculated using MAFFT v 7.310 (99,100) and the e-ins-i algorithm [options: --genafpair --maxiterate 1000 --seed] with the central part of the alignment available from NCBI CDD of the 100 most diverse sequences as seed. A consensus alignment was calculated using t-coffee v 11.00.8cbe486 (101) and used to construct a phylogenetic tree in FastTree v 2.1.10 (102) [options: ‐wag ‐gamma ‐slow ‐seed ‐notop ‐log] using the WAG-CAT model (103).

### Multiple sequence alignment, small sets

Three smaller sets were also aligned using MAFFT v 7.310 (99,100) and the l-ins-i algorithm [options: --localpair --maxiterate 1000]: Sir2 [Sir2Tm, Sir2La, F0THZ6 (*L. amylovorus*), A0A133PBX8 (*L. gasseri*), Q74LU0 (*L. johnsonii*), G9ZKM4 (*L. parafarraginis*), and CobB (*L. plantarum*)], SirTM [spySirTM, sauSirTM, LBA1649, F0TGP4 (*L. amylovorus*), D1YLJ9 (*L. gasseri*), A0A137PK42 (*L. johnsonii*)], and a set including the human sirtuins (hSIRT1-7, Sir2La, Sir2Tm, LBA1649, spySirTM, and sauSirTM). The latter set was used to calculate identities and similarities at imed.med.ucm.es/Tools/sias.html (similarity groups: FYW, RKH, DE, ST, NQ, and VILMA) (Table S2).

### BLAST search, Sir2La-like proteins

A BLASTP search of lactobacillus proteins in the NCBI database (blast.ncbi.nlm.nih.gov) using the Sir2 set discussed above as probes (E-value cut-off = 10^−6^) and removing closely related intraspecies sequences identified 269 sequences, spanning 156 different bacteria (Table S4).

### BLAST search and multiple sequence alignment, sirTM/LBA1649-like proteins

A BLASTP search of lactobacillus proteins in the NCBI database (blast.ncbi.nlm.nih.gov) using the sirTM set discussed above as probes identified 239 sequences, spanning 53 different bacteria. 100 alignments of these proteins were calculated using MAFFT v 7.310 (99,100) and the l-ins-i algorithm [options: --localpair --maxiterate 1000] and a consensus alignment was calculated using t-coffee v 11.00.8cbe486 (101). Based on the three residues discussed in the text to be critical for SirTM function, three groups could be identified: SirTM-like [NQR, 75 sequences, 29 species], LBA1649-like [XFP (X=A/S/T/Y), 104 sequences, 27 species], a smaller group [NEX (X=A/P/T), 20 sequences, 11 species], and additionally sequences with one or more of the residues missing (45 proteins) could be identified but were discarded in the following analysis (Table S4).

### Protein expression and purification

The gene encoding Sir2La was amplified by PCR from genomic DNA of *L. acidophilus* NCFM (generous gift of Dr. Morten Ejby) and subcloned into the pET-28a(+) vector using NdeI and BamHI restriction enzymes. The construct was transformed into DH5alpha cells and verified by sequencing (Eurofins Genomics and GATC Biotech), followed by transformation into BL21(DE3) competent *E. coli* cells. Cells were cultured in LB medium at 37 °C until OD_600_ 0.6–0.9, induced with 0.1 mM IPTG for gene expression, incubated at 20 °C for 16 h and harvested (14,000 *g*, 4 °C, 20 min). The cells were lysed by homogenization (Stanstead Pressure Cell Homogenizer, SPCH-10) in equilibration buffer A (25 mM HEPES, pH 7.4, 500 mM NaCl). The resulting lysate was clarified by centrifugation (14,000 *g*, 4 °C, 20 min) and loaded onto a 1 mL HisTrap HP (Äkta purifier, GE Healthcare), washed with equilibration buffer A (20 mL) and eluted by equilibration buffer A supplemented with 500 mM imidazole. Following analysis by SDS-PAGE, the appropriate fractions were combined and concentrated (10 kDa cut-off, Amicon Spinfilter). The buffer was exchanged (10 kDa cut-off, Amicon Spinfilter) to equilibration buffer B (50 mM sodium acetate, pH 5, 25 mM NaCl) and the resulting solution was loaded on a 1 mL HiTrap Capto Q/S column (Äkta purifier, GE Healthcare), washed with equilibration buffer B (approx. 20 column volumes), and eluted using a linear gradient of equilibration buffer B (approx. 40 column volumes) supplemented with sodium chloride to a final concentration of 1 M. After analysis by SDS-PAGE, the appropriate fractions were concentrated (as above). Protein concentration of the resulting solution was determined spectrophotometrically at 280 nm using theoretical molar extinction coefficients of 19.4·10^−3^ M^−1^cm^−1^ and 46.0·10^−3^ M ^−1^cm^−1^ for Sir2La and LBA1469, respectively (http://web.expasy.org/protparam) to 2.7-5.1 mg/mL (See the Supporting Information for full-length amino acid sequences of the produced proteins).

### Chemical synthesis

Fluorogenic substrates from the series **1** and substrates **2c** and **2d** were synthesized using protocols similar to previous reports (see the Supporting Information for full experimental details). The remaining substrates from series **2** and substrates from series **3**-**5** were reported earlier (44,60,61,70,104,105,106) and synthesized using a combination of solution-phase and solid-phase peptide synthesis (see the Supporting Information for references). Compound **6** has previously been reported (65) and compound **7** was synthesized from bis(1-benzotriazolyl)methanethione, methyl amine, and Cbz-Lys-Trp-NH*i* Pr·TFA using a protocol reported in the same paper (see the Supporting Information for full experimental details).

### Fluorescence-based deacylase assays

All reactions were performed at 25 °C in black low binding 96-well microtiter plates (Corning half-area wells), with duplicate series in each assay, and each assay was performed at least twice. Control wells without enzyme were included in each plate. All reactions were performed in assay buffer (50 mM HEPES, 100 mM KCl, 200 μM TCEP, 0.001% Tween-20 (v/v), pH 7.4, 0.5 mg/ml BSA) with appropriate concentrations of substrates and inhibitors obtained by dilution from 10–250 mM stock solutions in either water or DMSO and an appropriate concentration of enzyme obtained by dilution of the stock obtained as described above. All plates were analyzed using a PerkinElmer Life Sciences Enspire plate reader with excitation at 360 nm and detecting emission at 460 nm. Fluorescence measurements (relative fluorescence units) were converted to [AMC] concentrations based on an [AMC]-fluorescence standard curve. All data analysis was performed using GraphPad Prism v 7.0c.

### End-point fluorophore-release assays. Fluorogenic sirtuin substrate screening

The initial screening for substrate deacylation activity was performed with end-point fluorophore release by trypsin.(104,105,107) For a final volume of 25 μL/well, protein (final concentration, 250 nM), acyl substrate (50 μM), and either NAD^+^ or NADP^+^ (500 μM) was incubated (60 min), then a solution of trypsin and nicotinamide (25 μL, 5.0 mg/mL and 4 mM; final concentration of 2.5 mg/mL and 2 mM) was added, and the assay development was allowed to proceed for 90 min at 25 °C before fluorescence analysis. Co-substrate selectivity was determined using the same protocol. *End-point concentration-response (inhibitor) assays*—End-point inhibition assays (concentration-response) were performed in a final volume of 25 μL/well, where Sir2La (250 nM) and inhibitor (nicotinamide, ADPR, suramin, AGK2, SirReal2, sirtinol, **6** or **7**, 5-fold dilution series) was incubated with substrate (**2b**-**d** at *K*_M_ [**2b** 21 μM, **2c** 14 μM, and **2d** 15 μM]) and NAD^+^ (500 μM) at 25 °C for 60 min, then a solution of trypsin and nicotinamide (25 μL, 0.4 mg/mL and 4 mM; final concentration of 0.2 mg/mL and 2 mM) was added, and the assay development was allowed to proceed for 15 min at 25 °C before fluorescence analysis. For inhibitors, where residual activities below 50% were obtained at the highest tested concentrations, the data were fitted to the concentration-response equation to obtain IC50-values (μM).

### Continuous fluorophore-release assays. Michaelis-Menten plots and progress assays

Rate experiments for determination of kinetic parameters were performed in a final volume of 50 μL/well, where Sir2La (250 nM) and substrate (**2b-d**, 2-fold dilution series, 200-0.39 μM) were incubated with NAD^+^ (500 μM) and trypsin (10 ng/μL). In situ fluorophore release was monitored immediately by fluorescence readings recorded every 30 s for 60 min at 25 °C, to obtain initial rates *v*_0_ (nM s^−1^) for each concentration. The data were fitted to the Michaelis–Menten equation to afford *K*_M_ (μM) and *k*_cat_ (s^−1^) values. *Bisubstrate kinetics—* Rate experiments for determination of bisubstrate kinetic parameters were performed using the same protocol but employing substrate (**2c**, 2-fold dilution series, 200-1.56 μM) and NAD^+^ (2-fold dilution series, 1000-63 μM). The data were fitted to the global equation for a two-substrate rapid equilibrium random mechanism (Eq. 1), the global equation for a two-substrate steady-state ordered mechanism (Eq. 2, which has the same form as Eq. 1), or the global equation for a two-substrate rapid equilibrium ordered mechanism (Eq. 3) to afford dissociation and rate constants (Fig. 3a and Table 3). Evaluation of the model-fits was performed by replotting values obtained by individual fits to Eq. 3 and 4 for random and steady-state ordered mechanisms (Fig. 3b), or to Eq. 6–9 for the rapid equilibrium ordered mechanism assuming (for the resulting fits, see Supporting Figure S2).

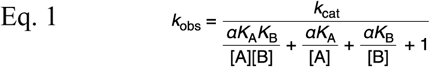

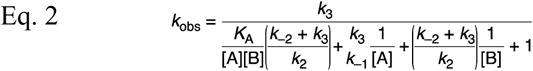

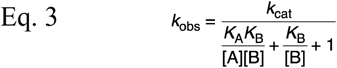

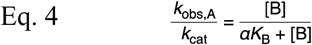

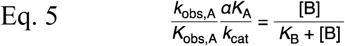

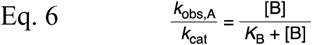

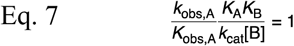

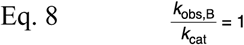

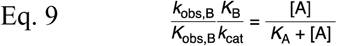

### Continuous concentration-response (inhibitor) assays

Rate experiments for determination of inhibition constants were performed using the same protocol, and using inhibitor (2-fold dilution series, nicotinamide 5000-9.8 μM; **6**, 300-0.59 μM), substrate (at substrate *K*_M_ [**2b** 21 μM and **2c** 14 μM]), NAD^+^ (500 μM) and trypsin (10 ng/μL). The data were then fitted to the concentration-response equation to obtain IC_50_-values (μM).

### HPLC-MS-based assays

For HPLC-MS-based assays, the same assay buffer was used, but with omission of BSA. Inhibition assays (concentration–response) were performed in a final volume of 25 μL/well, where Sir2La (250 nM) and NADH or NADPH (seven-point dilution series, 10– 0.01 mM) was incubated with substrate (**2b** or **2c**, 100 μM) and NAD^+^ (500 μM) at 25 °C for 60 min, followed by addition of MeOH/HCOOH (96:4 (v/v), 25 μL). The samples were analyzed by HPLC-MS on a Waters Acquity ultra-HPLC-MS system equipped with a diode array detector. A gradient with eluent I (0.1% HCOOH in water (v/v)) and eluent II (0.1% HCOOH in acetonitrile (v/v)) rising linearly 0–10% during t = 0.00–2.10 min followed by 10–60% of eluent II during t = 2.10–4.30 min was applied at a flow rate of 0.6 ml/min. The obtained chromatograms at 326 nm (where the coumarin moiety has its absorption maximum) were used to determine reaction progression, by determining area under the curve of the relevant peaks and the obtained mass spectra used to determine formation of the desired deacylated product using Waters MassLynx. For inhibitors, where residual activities below 50% were obtained at the highest tested concentrations, the data were fitted to the concentration-response equation to obtain IC_50_-values (μM).

## Acknowledgements

Karina Jansen is thanked for expert general assistance in the laboratory and Jacob Nedergaard Pedersen for help with protein production. We are grateful to Morten Ejby for the *L. acidophilus* NCFM genomic material used in this study and to Per Hägglund as well as to Ditte Welner for fruitful discussions.

## Author contributions

Conceptualization, Methodology, writing – review & editing, supervision, B.S., C.A.O., and A.S.M.; Investigation, S.V.O., N.R., C.A.O. and A.S.M.; Writing – original draft, S.V.O. and A.S.M.; Funding acquisition, B.S. and C.A.O.

## FOOTNOTES

This article contains Supporting Tables S1–S5, Figures S1–S3, amino acid sequences for produced proteins, synthetic methods, and ^1^H and ^13^C NMR-spectra for novel compounds.

This work was supported by the Technical University of Denmark, the Danish Research Foundation and Nordic Bioscience A/S (joint PhD fellowship to S.V.O.), the University of Copenhagen (PhD fellowship to N.R.), the Carlsberg Foundation (2011-01-0169, 2013-01-0333, and CF15-011; C.A.O.), the Novo Nordisk Foundation (NNF15OC0017334; C.A.O.), and the European Research Council (ERC-CoG-725172-SIRFUNCT; C.A.O.)

The abbreviations used are: ADPR, adenosine diphosphate ribose; AMC, 7-amino-4-methylcoumarin; ART, adenosine diphosphate-ribosyl transferase; cADPRS, cyclic adenosine diphosphate synthetase; DLAT, dihydrolipoyllysine acetyltransferase; GcvH-L, Glycine cleavage system H-like; H3, histone 3; H4, histone 4; HDAC, histone deacetylase; IPTG, isopropyl *β*-D-1-thiogalactopyranoside; Kac, *ε-N-*acetyllysine; Kbut, *ε-N-*butyryllysine; Kpro, *ε-N-*-propionyllysine; *lam, L. amylovorus; lga, L. gasseri; ljo, L. johnsonii, lpl, L. plantarum; lpa, L. parafarringis;* LplA2, lipoate protein ligase A; NR, nicotinamide riboside; OAcADPR, 2′-O-acetyl adenosine diphosphate ribose; OPrADPR, 2′-O-propionyl adenosine diphosphate ribose; p53, tumor suppressor p53; PARP, poly(adenosine diphosphate ribose) polymerases; PTM, post-translational modification; Sir2, silent information regulator 2; SirTM, class M sirtuin; TCEP, tris(carboxyethyl)phosphine.

The following database entries are used in the text: From the NCBI conserved domain database: The Sir2 superfamily NCBI accession number cd00296, Class U from Gram-positive bacterial species and fusobacteria NCBI accession number cd01411, and Class U with Sir2Af2 homology NCBI accession number cd01413; Sirtuin sequence motifs used for identifying sirtuin domains can be accessed at eggNOG: COG0846, Pfam: pf02146, and PROSITE: ps50305; The amino acid sequences of proteins disscussed can be accessed through the UniProt database: Sir2La/LBA0117 under UniProt Q5FMQ6, LBA1649 under UniProt Q5FIL3, spySirTM under UniProt P0DN71, and sauSirTM under UniProt A0A0H3JQ59. See the Supporting Information for further protein/UniProt references. The atomic coordinates for crystal structures discussed are available from the PDB database: Sir2Tm in complex with an acetyllysine peptide under PDB ID 2h2d, Sir2Tm in complex with a propionyllysine peptide under PDB ID 3pdh, and Sir2Af2 in complex with a myristoyllysine peptide under PDB ID 4ywj.

### ORCiDs

Sita V. Olesen: –

Nima Rajabi: 0000-0002-9509-7540

Birte Svensson: 0000-0002-2993-8196

Christian A. Olsen: 0000-0002-2953-8942

Andreas S. Madsen: 0000-0001-7283-2090

